# Saccades orchestrate intraocular glucose to shape visual responses in birds

**DOI:** 10.1101/2025.04.17.649318

**Authors:** Xi Xu, Tong Xiao, Yibing Chen, Lijuan Song, Chen Wu, Qian Wang, Tao Zhang, Yan Yang

**Affiliations:** State Key Laboratory of Cognitive Science and Mental Health, Institute of Biophysics, Chinese Academy of Sciences, China; University of Chinese Academy of Sciences, China; Sino-Danish Center for Education and Research, University of Chinese Academy of Sciences, China; State Key Laboratory of Cognitive Science and Mental Health, Institute of Psychology, Chinese Academy of Sciences, China

**Keywords:** Oscillatory saccades, Birds, Retinal metabolism, Intraocular glucose, Visual responses

## Abstract

Birds exhibit remarkable vision ^1–4^ despite lacking the typical network of blood vessels in their eyes ^5–7^. The characteristic poses a long-standing question about how avian retinas fulfill high energy demands necessary for sight. Here we show that natural, rhythmic eye movements, known as oscillatory saccades, orchestrate intraocular metabolic dynamics and attention-guided visual processing in pigeons. In a series of integrated experiments, we monitored eye movements along with glucose levels in the eye and neuronal activity in key brain regions receiving direct input from the retina. We found that these saccades generate fluctuations in intraocular glucose concentrations, closely linked to changes in neuronal visual responses over timescales of seconds to minutes. Moreover, pharmacological manipulations that altered glucose availability and eliminated these oscillatory saccades resulted in corresponding shifts in neuronal responses, demonstrating a causal linkage between oscillatory saccades, metabolic regulation, and visual processing. These findings reveal a mechanism by which birds actively evoked saccades during attention-driven information gathering, propelling retinal metabolism and facilitating their vision in the absence of retinal vasculature. This study underscores the interplay among eye movements, metabolic regulation, and high-level visual performance, suggesting broader implications for how eye movements contribute to retinal health, attention, and visual function across species.

## Main

Birds present a fascinating paradox in the animal kingdom: despite having higher metabolic demands ^8^, larger eyes ^9,10^, superior visual acuity ^1–4^, and thicker retinas than mammals ^11–13^, they entirely lack retinal blood vessels ^5–7^ that are essential for nutrient delivery in primates ^14–16^. This raises an intriguing enigma: how do avian eyes fulfill the extensive metabolic needs of thick, high-acuity retinas in the absence of direct vascular support? Solving this puzzle requires uncovering the sophisticated strategies that birds have developed to sustain retinal health and visual performance without conventional vascularization. One potential mechanism centers on *oscillatory saccades*—rapid, repetitive eye movements characterized by small, rhythmic shifts that include cyclotorsional rotations around the eye’s visual axis (Fig. 1a and Extended Data Fig. 1a,b) ^17,18^. Composing approximately 10% to 25% of total viewing time, these saccades have been recognized for stabilizing the visual scene and optimizing environmental searching ^19–23^. Emerging evidence, however, suggests that oscillatory saccades may also facilitate intraocular nutrient transport ^24^, by acting as mechanical “agitators”. Through repetitive motion, these saccades could propel glucose and other nutrients from the pecten oculi—a comb-like, pigmented vascular structure unique to birds (Extended Data Fig. 1c) ^10,25^—toward the retina, ensuring metabolic support to retinal neurons.

**Figure 1.**
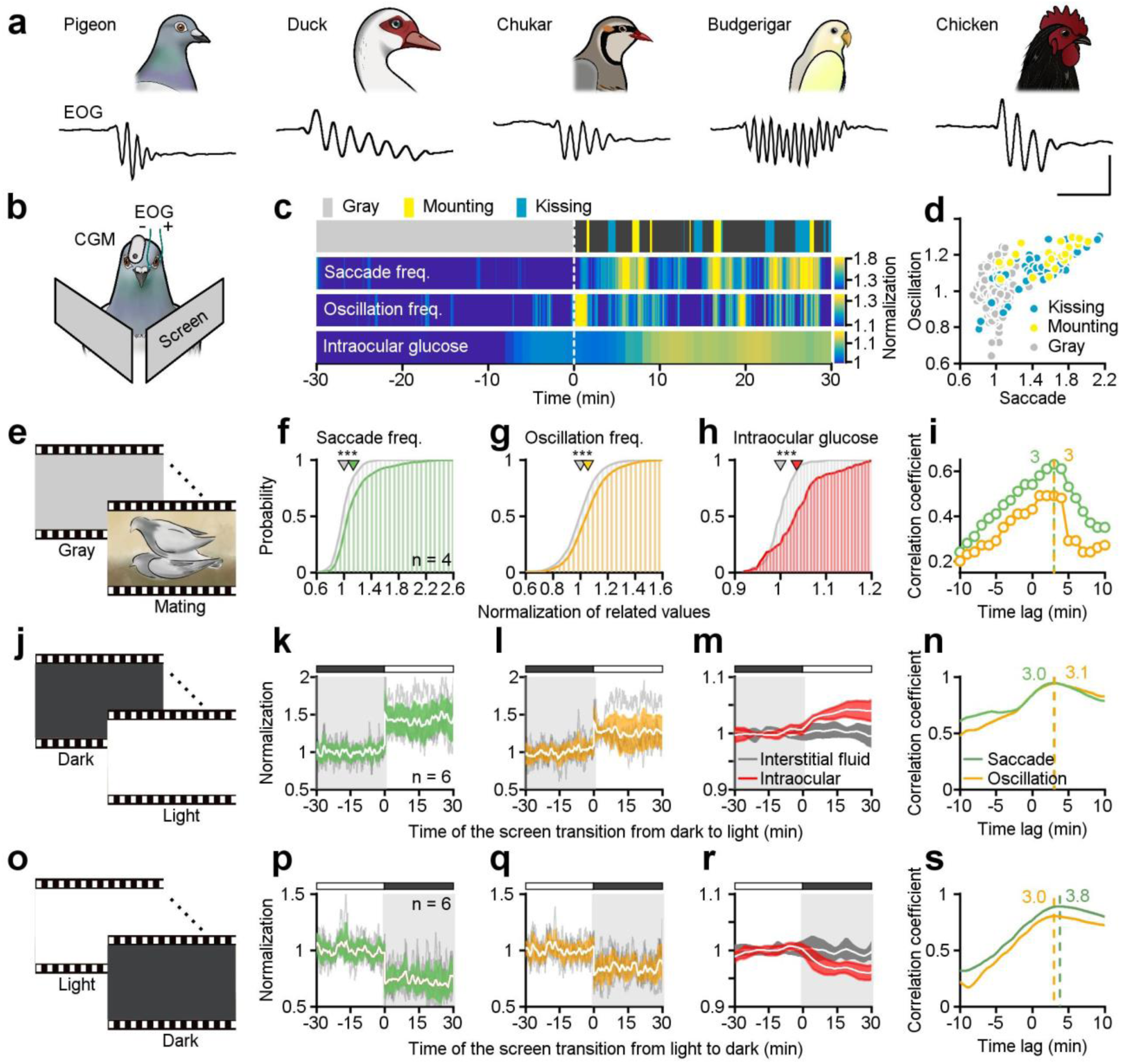
Oscillatory saccades driven by visual stimuli are correlated with intraocular glucose levels. **a,** Example electrooculography recordings of oscillatory saccades from five avian species: pigeon, duck, chukar, budgerigar, and chicken. Scale bars (left to right): 150 ms; 0.63 mV, 0.25 mV, 0.20 mV, 0.28 mV and 0.25 mV. **b,** Schematic drawing of the intraocular glucose monitoring (CGM) and eye movement (EOG) recording systems in awake-behaving pigeons viewing bilateral visual displays. **c,** An example pigeon exhibited increased saccadic eye movements and intraocular glucose levels while viewing 30-minute conspecific social mating videos. **d,** Both saccade frequency and oscillation frequency rose significantly during scenes involving kissing and mounting, as demonstrated by data normalized to gray screen baseline. **e,** Schematic diagram illustrating pigeon social mating videos. **f-h,** Cumulative distribution functions of saccade frequency (**f**), oscillation frequency (**g**), and intraocular glucose levels (**h**) during gray screen and mating-video viewing across four individuals. **i,** Correlation coefficients between intraocular glucose levels and both saccade frequency (green line) and oscillation frequency (yellow line) at different time lags throughout the mating video (Methods, Extended Data Fig. 3). **j,** Schematic diagram illustrating the dark-to-light screen transition. **k-m,** Normalized traces of saccade frequency (**k**), oscillation frequency (**l**), and glucose levels (**m**) during the transition from dark to light screens. The white lines with color shading represent the mean ± SD (n = 6 pigeons), and the gray lines indicate data from individual animals. **n,** Cross-correlation between saccade frequency and oscillation frequency with intraocular glucose levels dynamics (Methods). The dashed lines indicate the time lag with the highest correlation. **o-s,** Same as (**j-n**), but for the transition from light to dark screens. \*\*\**P* < 0.001, two-sided Wilcoxon rank-sum test.

These striking properties of avian eyes led us to hypothesize that their interplay may create a dynamic system for visual processing, driven by three interconnected mechanisms. First, oscillatory saccades are initiated for acquiring interesting information and sustaining retinal nutrient levels. Second, these eye movements facilitate the transport of metabolic substances, such as glucose ^26^, from the pecten oculi into the retina. Third, the resulting nutrient fluxes, especially the fluctuations in glucose levels, may dynamically modulate retinal function ^27^. Despite the compelling nature of these hypotheses, they have rarely been investigated as a cohesive entity, leaving critical gaps in our understanding of how avian eyes function without direct vascularization. Previous work has approached oscillatory saccade as an isolated, static event, by measuring the leakage of dye from the pecten following a single saccade ^24^. The challenge remains to establish cause-and-effect relationships among oscillatory saccades, dynamics of intraocular metabolism, and changes in neuronal responses during natural viewing behavior.

Here, we address this challenge using an integrated series of experiments designed to test whether —and how—oscillatory saccades drive fluctuations in intraocular glucose, and in turn, modulate visual responses in awake, behaving pigeons. We monitored eye movements and intraocular glucose levels while recording from neurons in three critical retinorecipient nuclei—key structures receiving direct input from retinal ganglion cells and essential for avian visual processing ^23,28–35^. We found that oscillatory saccades drive fluctuations in intraocular glucose, and dynamically correlate with changes in neuronal responses over timescales. Furthermore, pharmacological manipulation of both intraocular glucose and oscillatory saccades resulted in corresponding shifts in neuronal visual responses, establishing causal links. We report strong evidence showing how oscillatory saccades and glucose dynamics interact to shape visual processing in birds.

### Oscillatory saccades elicited by visual inputs are linked to intraocular glucose perfusion

Birds, like primates, exhibit selective attention through saccadic eye movements that prioritize salient visual stimuli ^21,36–39^. However, unlike primates, birds produce oscillatory saccades spanning tens to hundreds of milliseconds per event, with each oscillatory cycle lasting about 20 to 70 ms, depending on species and individual variation (Fig. 1a and Extended Data Fig. 1d-i and Supplementary Videos 1-5). These oscillatory saccades serve for efficient environmental scanning and attention-driven visual processing ^36,40^. Previous evidence indicates that they may also promote intraocular glucose perfusion through mechanical agitation of ocular fluids ^24^. To investigate how visual-guided saccades influence metabolic regulation, we simultaneously monitored eye movements (via electrooculography, EOG) and intraocular glucose concentrations (via clinical-grade Continuous Glucose Monitoring, CGM) in awake-behaving pigeons (Fig. 1b). The CGM system provided continuous, real-time glucose assessments (Extended Data Fig. 2a) with accuracy comparable to that of standard glucometers (Extended Data Fig. 2b), and did not significantly alter saccade (*P* = 0.20) or oscillation frequencies (*P* = 0.10).

When presented with 30-minute conspecific social mating videos preceded by a gray screen (Fig. 1e), pigeons displayed robust increases in both saccadic eye movements and intraocular glucose levels. Notably, saccade and oscillation frequencies rose significantly during scenes of kissing and mounting (an example animal in Fig. 1c,d, *P_saccade_* = 2.85 × 10^−32^, *P_oscillation_* = 7.68 × 10^−23^). Across four pigeons, saccade frequency, oscillation frequency, and intraocular glucose concentration increased by 16.08 ± 8.59%, 6.74 ± 2.30%, and 4.94 ± 1.55%, respectively, relative to a gray-screen baseline (*P_saccade_* = 4.58 × 10^−15^, *P_oscillation_* = 9.77 × 10^−7^, *P_glucose_* = 5.62 × 10^−24^; Fig. 1f-h). Correlation analysis revealed a temporal lag of approximately 3 minutes between glucose levels and both saccade and oscillation frequencies during the social visual task (Fig. 1i and Extended Data Fig. 3a,b). These findings indicate that attention-driven oscillatory saccades in birds enhance glucose mobilization from the pecten oculi to the retina, thereby boosting nutrient supply during social information gathering.

To determine whether non-social visual stimuli elicit comparable effects, we employed 2-hour screen-brightness cycles (1 hour dark/1 hour light; Fig. 1j). Within seconds of each dark-to-light transition, saccade frequency increased by 41.66 ± 6.95%, oscillation frequency by 25.62 ± 6.32%, and intraocular glucose levels by 3.35 ± 0.34% (*P_saccade_* = 0.00, *P_oscillation_* = 0.00, *P_glucose_* = 8.04 × 10^−18^, n = 6 pigeons; Fig. 1k-m). Conversely, light-to-dark transitions prompted reductions in saccade frequency (27.26 ± 3.14%), oscillation frequency (16.43 ± 2.60%), and intraocular glucose (2.88 ± 0.58%) (*P_saccade_* = 0.00, *P_oscillation_* = 0.00, *P_glucose_* = 8.46 × 10^−11^; Fig. 1o-r). Simultaneous measurements of interstitial glucose ruled out systemic stress responses (dark-to-light: *P* = 0.08; light-to-dark: *P* = 0.41). Further, lag correlation analysis revealed that intraocular glucose changes followed fluctuations in saccade and oscillation frequencies by approximately 3-4 minutes (Pearson correlation coefficient; Fig. 1n,s), indicating a similar temporal delay between mechanical agitation of intraocular fluids and enhanced nutrient delivery to retinal tissues. These results demonstrate that avian oscillatory saccades actively propel glucose and other nutrients towards the retina, providing a critical metabolic boost for visual processing with a temporal lag of several minutes.

### Oscillatory saccades dynamically correlate with retinorecipient neuronal responses across timescales ranging from seconds to minutes

To explore whether oscillatory saccades modulate neuronal visual responses, we investigated three crucial retinorecipient nuclei that receive direct retinal input and subserve distinct functions within the avian visual pathways. Using a large-field still grating that nearly filled the pigeons’ right visual field, we simultaneously monitored eye movements and conducted extracellular recordings in these nuclei. To evoke robust visual responses, the grating periodically moved at 8°/s for 300 ms at random intervals (6 to 10 seconds), with motion direction optimized for each neuron’s preference (Fig. 2a). These stimuli were delivered during fixation periods between oscillatory saccades, enabling systematic probing of neuronal excitability to visual inputs while minimizing interference from eye movements-related artifacts (Extended Data Fig. 4). In total, we recorded 127 neurons: 57 in the optic tectum (OT) of the tectofugal pathway, 20 in the nucleus opticus principalis thalami (nOPT) of the thalamofugal pathway, and 50 in the pretectal nucleus lentiformis mesencephali (nLM) of the accessory optic system (Fig. 2b).

**Figure 2.**
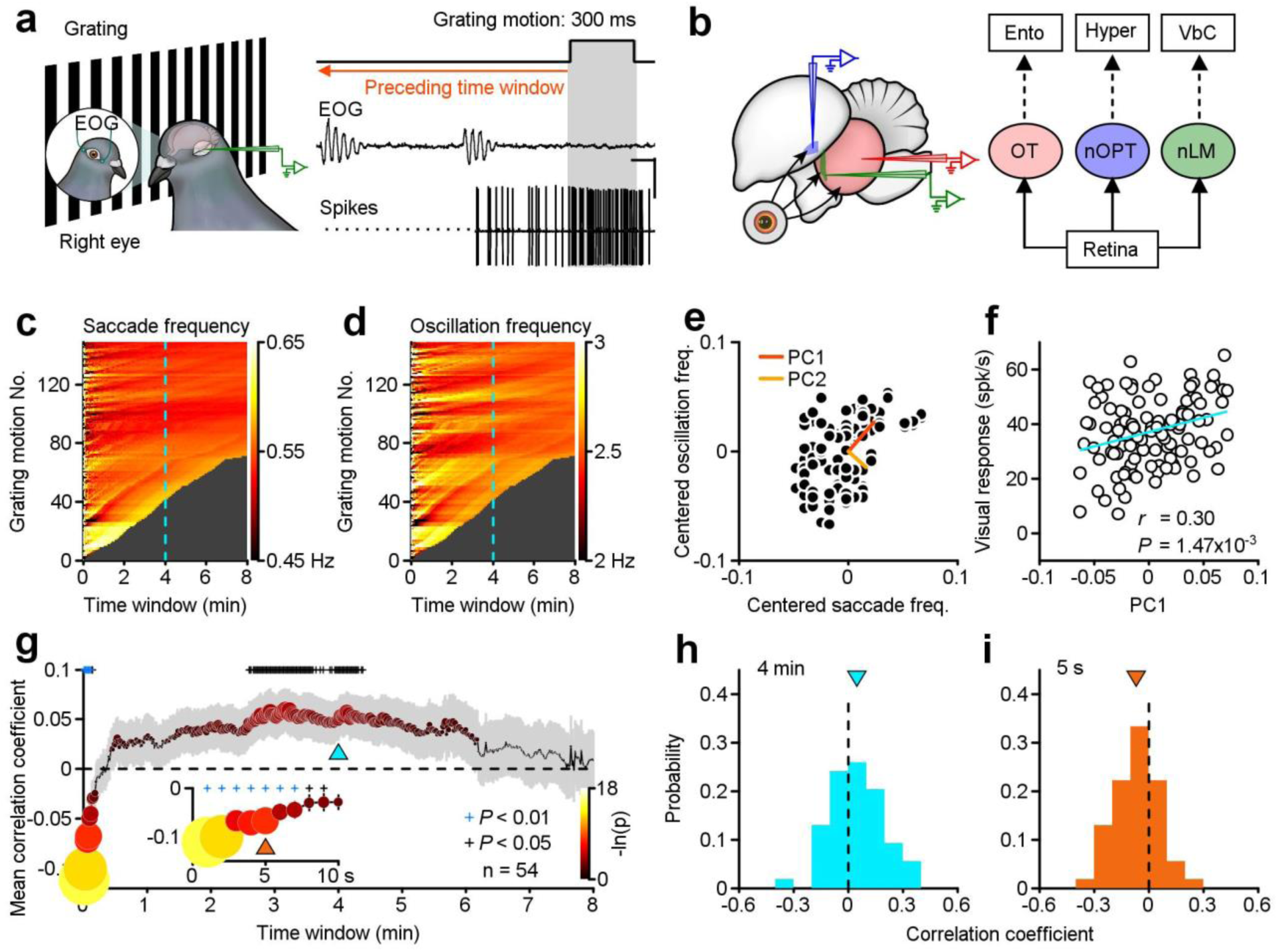
Oscillatory saccades dynamically correlate with visual responses in retinorecipient neurons across both second- and minute-timescales. **a,** Schematic of simultaneous recordings of eye movements and neuronal activity in pigeons during exposure to grating motion. Right: Representative raw traces of EOG and single-neuron activity, with gray shading marking grating motion. The reddish-orange arrow points to the preceding time window analyzed for oscillatory saccades prior to grating motion. Scale bars: 100 ms; 0.33 mV (EOG); 0.5 mV (spikes). **b,** Recordings targeted brain regions receiving direct retinal projections: OT, nOPT and nLM. The right panel illustrates the three principal avian visual pathways. Ento, Entopallium; Hyper, Hyperpallium; Vbc, Vestibulocerebellum. **c,d,** Saccade frequency (**c**) and oscillation frequency (**d**) within preceding time windows (1 second to 8 minutes), aligned with grating motion for an example neuron. Dashed sky-blue lines indicate a 4-minute time window. **e,** Principal component analysis of saccade and oscillation frequencies across the time window. **f,** Correlation between visual responses of the example neuron and PC1 scores derived from PCA in (**e**). **g,** Correlation coefficients between PC1 of oscillatory saccades and neuronal visual responses across preceding time windows of different durations. Symbol size and color reflect the degree of statistical significance, represented as −ln(p) (n = 54, two-sided Wilcoxon signed-rank test); gray shading represents 1 SEM. The inset highlights result for time windows ≤ 10 seconds. The sky-blue and orange upward triangles indicate the 4-minute and 5-second time windows, respectively. **h,i,** Population distribution of correlation coefficients across all 54 neurons for the 4-minutes (**h**) and 5-second (**i**) time windows.

We investigated how variability in visual responses of retinorecipient neurons related to preceding oscillatory saccades. To systematically assess this relationship, we analyzed eye movements occurring over various intervals prior to grating motions, ranging from 1 second to 8 minutes in 1-second increments (example neuron’s data illustrated in Fig. 2c,d). We utilized principal component analysis (PCA) to quantify combined effects of saccade and oscillation frequencies. Using a representative 4-minute window (dashed sky-blue lines), PCA decomposed eye movement data into principal components (Fig. 2e). Correlation analysis demonstrated a significant positive relationship between the first principal component (PC1) and corresponding neuronal visual responses (r = 0.30, P = 1.47 × 10^−3^, n = 108, Pearson correlation; Fig. 2f). This indicates that more frequent oscillatory saccades within the preceding period relate to enhanced neuronal activity.

For the population analysis, we focused on 54 out of 127 neurons that had at least 35 repetitions of the moving grating per time window, extending up to 8 minutes. Intriguingly, these retinorecipient neurons revealed both short- and longer-term correlations between their visual responses and preceding oscillatory saccade activity. On a minute-scale, positive correlations emerged approximately 3-4 minutes prior (n = 54, *P* < 0.05; Fig. 2g,h). Notably, this interval aligns with the lag observed between saccadic eye movements and intraocular glucose changes (Fig. 1). It suggests that oscillatory saccades could alter intraocular glucose delivery within this timeframe, thereby modulating retinal visual responses and influencing subsequent neuronal processing at higher visual regions. Conversely, on a second-scale, negative correlations were evident within the first 10 seconds, particularly pronounced during the initial 7 seconds (*P* < 0.01; Fig. 2g,i), indicating a short-term effect likely driven by the most recent one or two saccades.

To further examine the short-term effect in more detail, we examined how each neuron’s responses varied with the time elapsed since the latest oscillatory saccade. As an example, one neuron’s visual responses were sorted by these intervals spanning up to 6 seconds and binned into three stages with equal ranges (Fig. 3a), revealing a linear increase in firing as time from the prior saccade onset (*r* = 0.43, *P* = 1.92 × 10^−8^; Fig. 3b). Across these three stages, z-scored visual responses rose significantly (one-way ANOVA, *F_(2,155)_* = 13.13, *P* = 5.41 × 10^−6^; Fig. 3c). Population-level analysis across the three visual nuclei showed stronger visual responses in stage 3 than in stage 1 (OT: *P* = 2.71 × 10^−6^; nOPT: *P* = 5.11 × 10^−3^; nLM: *P* = 1.11 × 10^−4^; Fig. 3d-f), with the majority of neurons exhibiting this pattern (OT: 48/57, 84%; nOPT: 16/20, 80%; nLM: 36/50, 72%). Furthermore, visual responses were positively correlated with the time since the latest saccade (OT: *r* = 0.44, *P* = 1.44 × 10^−9^; nOPT: *r* = 0.47, *P* =1.75 × 10^−4^; nLM: *r* = 0.36, *P* = 6.05 × 10^−6^; Pearson correlation coefficient; Fig. 3g-i). The same effects persisted after regrouping intervals to ensure equal repeat counts (Extended Data Fig. 5).

**Figure 3.**
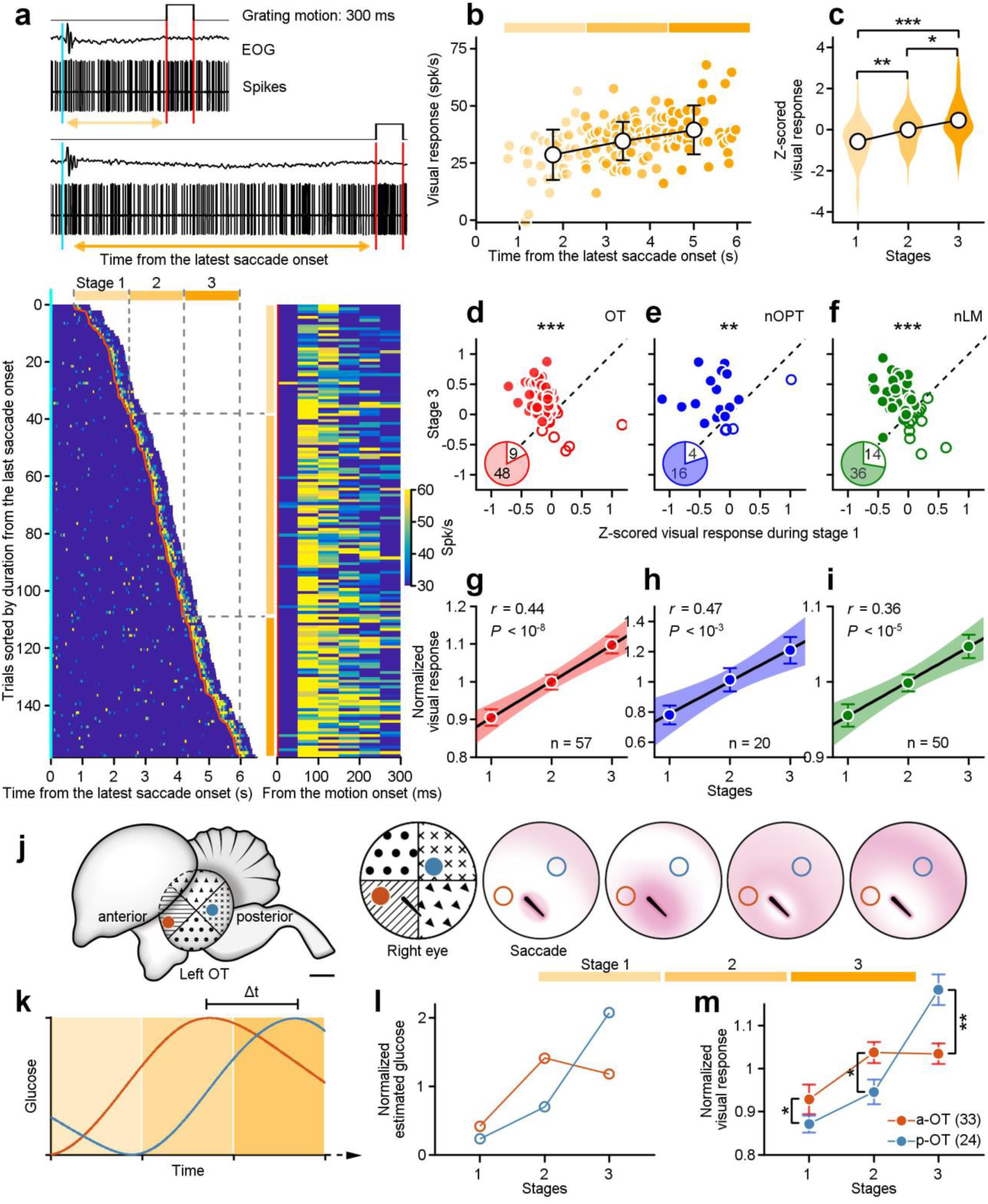
Dynamic changes in visual responses following the latest oscillatory saccade. **a,** Example traces of EOG and neuronal spiking activity in response to grating motion. Red lines mark motion onset/offset, while sky-blue lines indicate the latest saccade onset. Visual responses were categorized into three stages based on intervals from the saccade onset, represented by progressively darker yellow shades. Heatmaps depict responses from an example neuron, sorted by time since the latest saccade onset (left) and aligned to grating motion onset (right). **b-c,** Visual responses of the example neuron plotted against time since the prior saccade onset (**b**) and z-scored distributions across three stages (**c**) (one-way ANOVA, *F*_(2, 155)_ = 13.13, *P* = 5.41 × 10^−6^). Post hoc pairwise comparisons were adjusted using the Holm-Bonferroni (adjusted *P* < 0.02). **d-i,** Comparison of z-scored visual responses between stage 3 and stage 1 for neurons in the OT (**d**), nOPT (**e**), and nLM (**f**) regions, and their normalized visual responses across three stages (**g-i**). Black lines represent the linear regression, while shadings are the 95% confidence interval of fitting. **j-l,** Estimated glucose dynamics following a saccade differ between anterior OT (a-OT, red) and posterior OT (p-OT, blue) neurons due to the proximity of their visual receptive fields to the pecten^30,42,43^. Red and blue dots indicate the recording sites and their respective receptive fields. Temporal glucose diffusion patterns reveal a delay between a-OT and p-OT neurons (**k**), with differences across stages (**l**). **m,** Visual responses of a-OT and p-OT neurons showed significant differences during each stage. Data points and error bars represent mean ± SEM. * *P* <0.05, ** *P* <0.01, *** *P* <0.001, Two-sided Wilcoxon rank-sum test.

These second-scale effect may stem from the time required for glucose and other nutrients to diffuse across the retina after each oscillatory saccade. If this is the case, neurons whose receptive fields lie closer to the pecten should be modulated sooner, reflecting a gradient of glucose availability (Fig. 3j). To test this, we focused on OT neurons with receptive fields at different retinal locations. In the avian optic tectum or the mammalian superior colliculus, visual neurons receive topographically organized retinal projections ^41,42^. We recorded from OT neurons in two distinct regions: an anterior area (red spot, A 5.00-6.00, n = 33 neurons), whose receptive fields situated nearer the pecten, and a posterior area (blue spot, A 1.00-1.25, n = 24 neurons), located farther away ^30,42,43^. Consequently, following oscillatory saccades, glucose was estimated to travel shorter distances to reach anterior regions faster, producing distinct temporal patterns of nutrient availability across the retina for anterior and posterior OT neurons (red vs blue lines in Fig. 3k). Indeed, visual responses in both anterior and posterior OT neurons tracked these estimated glucose dynamics (Fig. 3l,m). Compared with posterior OT neurons, anterior OT neurons were significantly more active in the initial two stages and then became less active in stage 3 (*P_1_* = 0.01; *P_2_* = 0.04; *P_3_* = 1.86 × 10^−3^; Fig. 3m). These findings support a diffusion-based mechanism for the second-scale modulation of visual responses, governed in part by the spatial proximity of receptive fields to the pecten.

### Intraocular glucose modulation alters visual responses

To investigate how intraocular glucose regulates neuronal visual responses, we locally manipulated intraocular glucose concentrations in awake pigeons while simultaneously recording neuronal activity and monitoring eye movements. Initially, intraocular glucose levels were elevated via intravitreal microinjections of D-glucose (1 mol/L, 2 μL; see Methods), raising intraocular glucose from 15.36 ± 0.40 to 17.69 ± 0.53 mmol/L (mean ± SEM, n = 9, *P* = 3.91 × 10^−3^). Due to spatial limitations in our recording setup, we focused on retinorecipient neurons in the nLM and measured visual responses to 300 ms grating-motion stimuli (Fig. 4a). In an example neuron, visual responses increased from 23.70 ± 13.22 to 70.23 ± 17.70 spikes/s within 2 minutes post-injection (mean ± SD, *P* = 8.16 × 10^−5^; Fig. 4c,d). Across all nine examined nLM neurons, D-glucose administration significantly enhanced visual responses (*P* = 7.81 × 10^−3^; Fig. 4g). Control experiments with L-glucose and saline injections revealed no significant changes in visual responses on the example neuron (Fig. 4e,f) or across the population (L-glucose: *P* = 0.30, n = 9 neurons, Fig. 4h; saline: *P* = 0.58, n = 7 neurons, Fig. 4i). The order of these injections (D-glucose, L-glucose, and saline) was randomized for each examined neuron.

**Figure 4.**
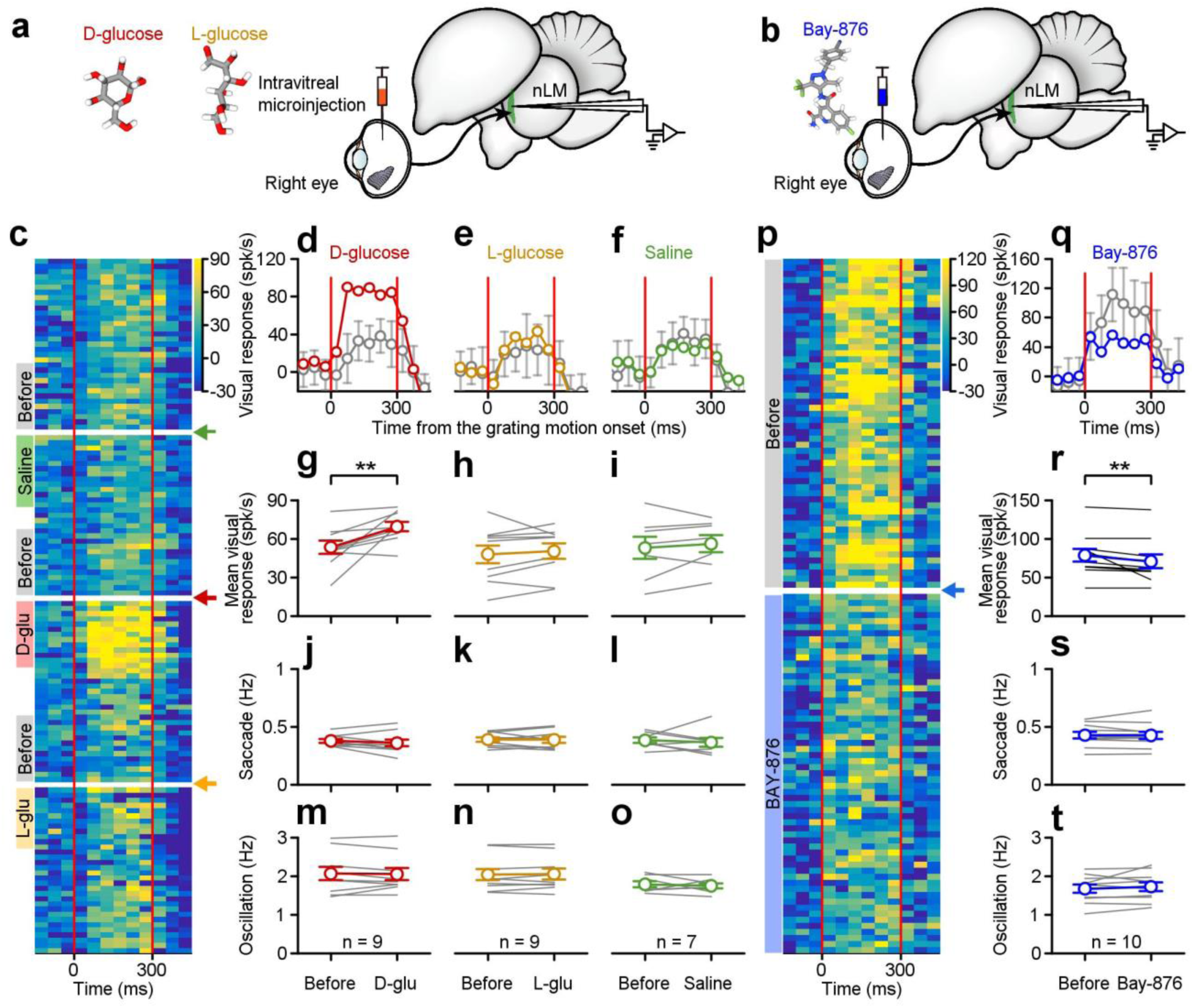
Bidirectional intraocular glucose manipulations effectively modulate visual responses. **a,b,** Pharmacological experimental setup illustrating intravitreal microinjections of D-glucose, L-glucose, saline (**a**) or Bay-876 (**b**). **c,** Heatmaps showing visual responses of an example neuron to grating motion, before and after intravitreal microinjections of saline, D-glucose, and L-glucose, in order. Colored arrows indicate the time of injections, and colored bands represent analysis windows (2 minutes). **d-i,** Visual responses before and after intravitreal microinjections of D-glucose (**d,g**), L-glucose (**e,h**), and saline (**f,i**) for the example neuron (**d,f**) and the neuron population (**g,i**). Shaded areas in (**d,f**) represent ±1 SD, while (**g,i**) depict individual neuron data (gray lines) and mean ± SEM (colored lines). **j-o,** Saccade frequencies (**j-i**) and oscillation frequencies (**m-o**) before and after injections of D-glucose (**j,m**), L-glucose (**k,n**), and saline (**l,o**). **p,** Heatmaps of visual responses from an example neuron before and after Bay-876 injection. The blue arrow marks the injection time, and colored bands represent the 10-minutes analysis windows before and after the injection. **q,r,** Visual responses before and after Bay-876 injection for an example neuron (**q**) and across the neuron population (**r**). **s,t,** Oscillatory saccade frequencies before and after Bay-876 injection. ** *P* < 0.01, Two-sided Wilcoxon signed-rank test.

We next explored the effects of decreasing intraocular glucose by inhibiting glucose transporter 1 (GluT-1; Fig. 4b), a primary glucose transporter highly expressed in the avian pecten oculi ^44,45^. Intravitreal administration of the GluT-1 inhibitor Bay-876 (100 μmol/L, 1 μL) ^46–48^ effectively reduced glucose concentrations from 15.82 ± 0.34 to 11.52 ± 1.31 mmol/L (mean ± SEM, *P* = 1.95 × 10^−3^, n = 10 pigeons). This decrease led to reduced visual responses, as illustrated by an example neuron, whose firing dropped from 85.67 ± 24.88 to 47.10 ± 16.21 spikes/s (mean ± SD, *P* = 1.25 × 10^−14^, Fig. 4p,q). Population analyses of 10 neurons confirmed that lowering intraocular glucose significantly diminished visual responses (*P* = 9.77 × 10^−3^; Fig. 4r).

None of these microinjections affected oscillatory saccades (Fig. 4j-o, and Fig. 4s,t) or neuronal spontaneous firing rates (*P* > 0.05). These findings demonstrate that pharmacological manipulating of intraocular glucose, either elevating or reducing it, robustly influences neuronal visual responses. They underscore the critical role of glucose in retinal function and its potential as a key modulator of sensory processing in visual systems.

### Muscimol-induced suppression of oscillatory saccades unmasks glucose-driven modulation of visual responses

To determine how oscillatory saccades modify neuronal visual responses through changes in glucose levels, we suppress these oscillatory eye movements by injecting the GABA_A_ receptor agonist muscimol into the raphe complex, key brainstem nuclei for saccade generation ^19,49,50^. Concurrently, we monitored intraocular glucose concentrations and neuronal activity in response to visual stimulation (Fig. 5a). After muscimol injection, both saccade and oscillation frequencies were significantly reduced (saccade frequency: pre-injection, 0.28 ± 0.04 Hz; post-injection, 0.02 ± 0.01 Hz, *P* = 4.19 × 10^−104^; oscillation frequency: pre-injection, 1.47 ± 0.27 Hz; post-injection, 0.04 ± 0.01 Hz, *P* = 1.59 × 10^−99^; mean ± SEM, n = 6 pigeons; Fig. 5b,c). These reductions were accompanied by a significant decrease in intraocular glucose, from 16.00 ± 2.01 to 11.45 ± 2.08 mmol/L (*P* = 3.36 × 10^−10^; Fig. 5d). Utilizing PCA on saccade and oscillation frequencies, along with lag correlation analysis between the PC1 and glucose fluctuations, we observed a 3.8-minute delay between reduced saccade activity and subsequent drops in intraocular glucose (Fig. 5e), mirroring the temporal lag found in previous experiments (Fig. 1).

**Figure 5.**
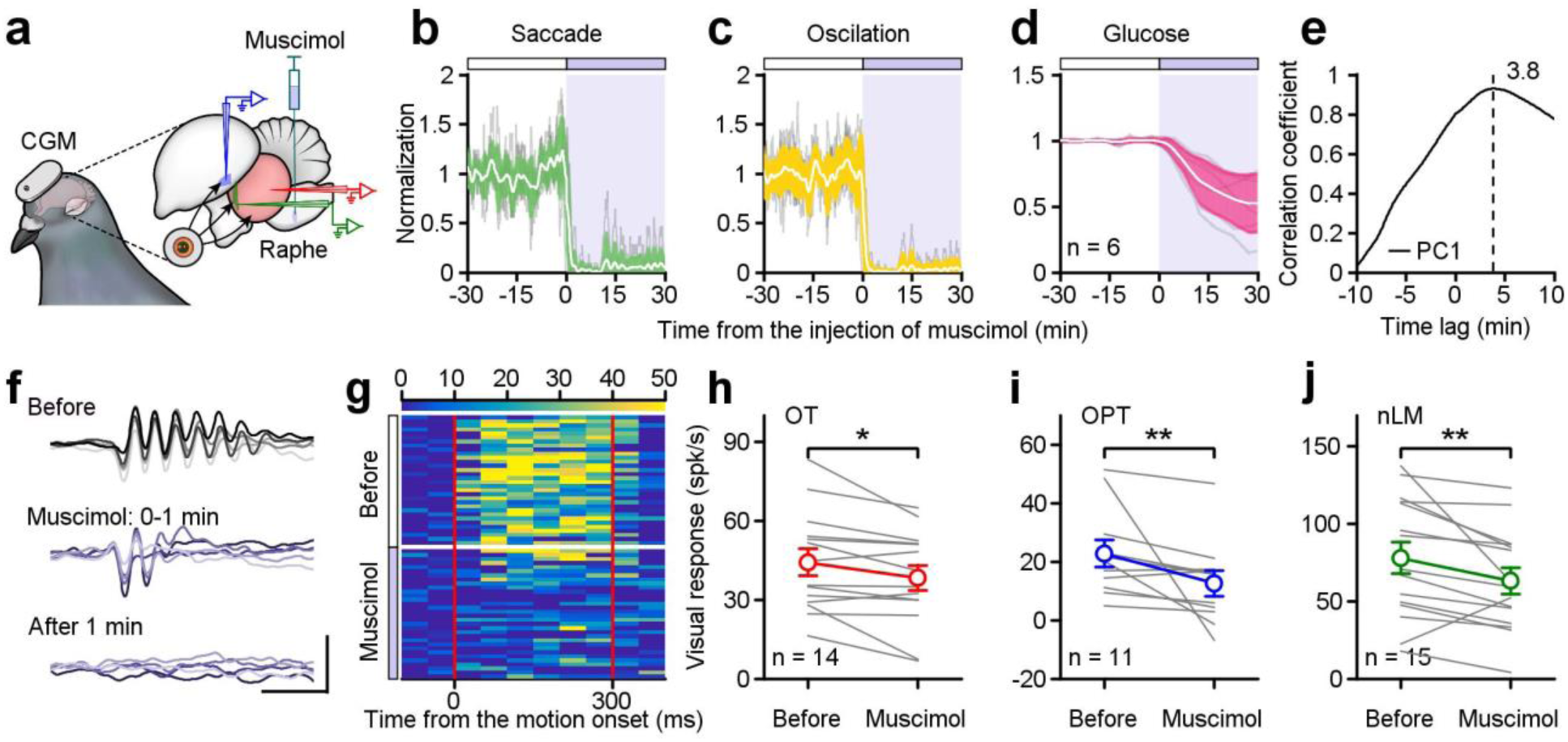
A causal link between oscillatory saccades, intraocular glucose regulation, and visual response modulation. **a,** Schematic of the experimental setup showing muscimol injections into the raphe complex to suppress oscillatory saccades during glucose monitoring and neuronal recordings in OT, nOPT and nLM. **b-d,** Normalized time-course changes in saccade frequency (**b**), oscillation frequency (**c**), and intraocular glucose levels (**d**) before and after muscimol injections. Thin gray lines represent individual animals, while white lines with colored areas show the population mean ± SD. **e,** Cross-correlation between intraocular glucose levels and eye movement dynamics, represented by PC1 of saccade and oscillation frequencies. The dashed line indicates the time lag with the strongest correlation. **f,g,** Saccadic eye movements (**f**) and visual responses of an example neuron (**g**) before and after muscimol injection. Scale bars: 100 ms; 0.2 mV. **h-j,** Comparison of visual responses to grating motion in the OT (**h**), nOPT (**i**), and nLM (**j**) within 10 minutes before and after muscimol injection. Gray lines represent individual neurons, while colored lines depict population means ± SEM. * *P* < 0.05, ** *P* < 0.01, Two-sided Wilcoxon signed-rank test.

Suppressing oscillatory saccades with muscimol significantly attenuated eye movements and visual responses, as demonstrated by an example neuron (Fig. 5f,g), and confirmed across all three examined nuclei (Fig. 5h-j). In the OT, neuronal activity to visual stimuli decreased from 44.10 ± 5.10 to 38.22 ± 4.81 spikes/s (mean ± SEM, *P* = 0.01, n = 14 neurons; Fig. 5h). Similarly, in the nOPT, visual responses dropped from 22.74 ± 4.54 to 12.59 ± 4.35 spikes/s (*P* = 6.84 × 10^−3^, n = 11 neurons; Fig. 5i), and in the nLM, declined from 77.89 ± 10.12 to 63.15 ± 8.40 spikes/s (*P* = 5.37 × 10^−3^, n = 15 neurons; Fig. 5j). In contrast, muscimol had no significant impact on spontaneous neuronal firing (*P_OT_* = 0.67, *P_nOPT_* = 0.08, *P_nLM_* = 0.12). Together, these results highlight the pivotal role of oscillatory saccades in regulating intraocular glucose and in modulating visual responses in retinorecipient neurons. The interplay between saccadic eye movements and metabolic dynamics offers mechanistic insights into how avian retinas meet high metabolic demands and maintain optimal visual processing.

## Discussion

Unlike primates, birds lack retinal blood vessels ^6,7^ and instead rely on alternative mechanisms to meet the metabolic demands of their thick, high-acuity retinas ^24,51^. Our findings reveal a sophisticated process wherein avian oscillatory saccades actively orchestrate retinal glucose fluctuations, and dynamically shape neuronal visual responses depending on the temporal context. Both minute- and second-scale modulations underscore the complexity of the relationship between eye movements and visual processing. Under natural viewing conditions, we observed an approximate 3-4 minute lag between attention-driven saccades and subsequent changes in intraocular glucose (Fig. 1), aligning closely with improvements in neuronal responses over the same timeframe (Fig. 2g,h). These findings suggest a beneficial minute-scale effect, wherein repetitive saccades gradually boost retinal responsiveness through saccade-driven glucose fluctuations. Such a mechanism likely ensures continuous metabolic support for optimal visual function in avian retinas lacking direct vascularization. It reflects long-term benefits of cumulative nutrient metabolism, linked with the state of eye movements over the longer term, and in turn, to enhance retinal function.

In contrast to the longer-term mechanism, our results also revealed negative correlations between eye movements and neuronal visual responses on the order of seconds (Fig. 2g,i). Specifically, visual responses increased progressively following individual saccades (Fig. 3), indicating a transient effect on visual processing. Previous research demonstrated that fluorescein dye disperses about 3 mm from the avian pecten within 2 seconds after a single saccade, whereas during prolonged fixation, dye diffusion was notably limited, less than 0.5 mm even after 120 seconds, highlighting the inefficiency of passive diffusion mechanisms ^24^. Thus, oscillatory saccades appear to actively drive nutrient perfusion from the pecten across the avian retina, achieving widespread metabolite distribution within seconds (Fig. 3j). Supporting this interpretation, our data confirmed that neuronal receptive fields nearer to the pecten experienced earlier enhancements than those farther away (Fig. 3j-m), consistent with a spatial gradient in nutrient availability occurring within seconds. This short-term, second-scale modulation likely served as an instant mechanism enabling swift metabolic perfusion post-saccade, transiently augmenting nutrient availability and enhancing subsequent retinal visual responses.

Our pharmacological manipulations provide a causal linkage between saccades, glucose dynamics and visual neuronal responses. Increasing intraocular glucose with D-glucose (but not the non-metabolizable L-glucose) ^52–54^, and decreasing it via the GluT-1 inhibitor Bay-876 ^46–48^, reliably altered neuronal visual responses (Fig. 4). These manipulations confirm that fluctuations in local glucose availability sensitively tuned retinal function, highlighting the critical role of precise metabolic regulation for sustaining high-acuity vision under varying environmental demands. Additionally, inactivation of the brainstem raphe complex via muscimol, a known gating hub for saccade generation ^19,50^, suppressed oscillatory saccades, subsequently lowering intraocular glucose levels and diminishing visual responses (Fig. 5). These findings reinforce the pivotal role of oscillatory saccades in ensuring adequate nutrient supply to retinal neurons.

It should be noted that oscillatory saccades in birds actively contribute to visual processing beyond their traditional role in environmental scanning. Our findings indicate that these saccades are not merely passive contributors, but function as “active drivers” of intraocular metabolic regulation and visual performance. During complex visual scenarios, such as social interactions, oscillatory saccades become more frequent and pronounced, acting as a mechanism for optimizing visual search efficiency (Fig. 1). These increased saccades are accompanied by elevated intraocular glucose levels, effectively meeting the heightened metabolic demands of retinal neurons engaged in processing complex visual information. Meanwhile, avian visual responses are not only sustained, but dynamically optimized to efficiently process salient and intricate visual stimuli by enhancing retinal energy availability. This interplay among saccadic eye movements, intraocular metabolic support, and neuronal responses underscores a sophisticated adaptive mechanism tailored to the high-acuity visual demands faced by birds. Specifically, the enhancement of oscillatory saccades during attention-intensive tasks ensures precise shaping of retinal function to cope effectively with visually challenging environments.

Our findings provide novel insights into the critical role of avian oscillatory saccades and extend their significance beyond basic visual scanning to metabolic and functional optimization. Moreover, they raise broader implications regarding the role of eye movements in maintaining retinal health, enhancing attentional performance, and optimizing visual function across species.

## Methods

### Animal preparation

We performed experiments on forty-seven awake-behaving adult pigeons (*Columba livia*, body weight 300-500 g; both sexes). Pigeons were randomly assigned to two sets of behavioral and physiological studies. In the first set, ten pigeons were utilized to investigate saccadic eye movements while viewing social mating videos or alternating light/dark screens; intraocular glucose levels were continuously monitored during these sessions. In the second set, the remaining thirty-seven were evaluated for neuronal visual responses to grating motion stimuli presented in a dimly lit room, associated with pharmacological manipulations. Among them, eighteen animals received muscimol injections targeting the raphe nucleus, nine animals were administrated intraocular injections of D-glucose, L-glucose and saline in a randomized order; and the remaining ten animals were given an intraocular injection of Bay-876, a potent and selective GluT-1 inhibitor. In the first set of experiments (Fig. 1 and Extended Data Fig. 1), we broadened the evolutionary scope by including four additional avian species—a budgerigar (*Melopsittacus undulatus*), a chukar partridge (*Alectoris chukar*), a chicken (*Gallus gallus domesticus*), and a muscovy duck (*Cairina moschata*)—which diverged tens of millions of years ago ^55^, to assess their oscillatory saccade patterns.

Each animal was anesthetized by injecting ketamine (30 mg/kg) and dexmedetomidine (0.125 mg/kg) into the pectoral muscles. The anesthetic status was monitored by examining their respiration patterns and reflex reactions to toe pinching ^19,23,56^. Pigeons were wrapped with soft straps and positioned into a stereotaxic apparatus. We made an incision on their scalp and attached a lightweight steel holder to their skull in order to secure head restraint in experiments. Utilizing stereotaxic coordinates of the pigeon brain ^57^, recording windows over the left tectum, the left telencephalon, as well as an injection window over the cerebellum were surgically exposed by a dental drill. The overlying dura mater remained intact. We also gently drilled a little fragment of the orbital bone (1 × 1 mm) located on the top of the right eye, for the purpose of interocular glucose examinations and interocular injections during experiments. Following the exposure of these locations, we sutured the scalp closed and prescribed erythromycin ointment (BAIJINGYU Pharmaceutical, Nanjing, China). Pigeons were returned to their individual recovery cages and received ibuprofen (20 mg/kg, Sino-American Tianjin SmithKline & French Laboratories, China) to alleviate postoperative pain for several days after surgery. Within 12-24 hours, the pigeons’ locomotion, feeding, and hydration behaviors had returned to their normal state.

During an experimental day, the pigeon was administered a light anesthetic containing ketamine (4 mg/kg), coated in a bag, and situated on a foam couch. The head was secured to the stereotaxic apparatus using a rod connected to a head holder. The wound margin and muscles were periodically infiltrated with lidocaine (Shanghaizhaohui Pharmaceutical, China). After the pigeon adapted to the restraint and remained unruffled on the couch, it was intramuscularly injected with ceftriaxone sodium (75 mg/kg) and the analgesic buprenorphine (0.02 mg/kg). During neuronal recordings, the left eye was covered while the right eye was kept open to visual stimuli. For monitoring eye movements in the budgerigar, chukar partridge, chicken and duck, animals underwent the same anesthesia and head fixation procedures as the pigeons. Procedures in this study were in strict accordance with the guidelines for the care and use of animals established by the Society for Neuroscience. All of the animals were handled according to protocols approved by the Institutional Animal Administration Committee at the Institute of Biophysics, Chinese Academy of Sciences (IBP-P-001(21) and IBP-P-001(24)).

### Visual conditions

The study introduced three types of visual stimuli to the animals, programmed using Psychtoolbox-3 (version 3.0.18) and executed within the MATLAB platform (R2016b, MathWorks Inc., USA). Mating videos and light/dark stimuli were presented on two 27-inch computer screens (Q27P2, AOC, China) positioned around the viewer’s head (Fig. 1b). In the video experiments, pigeons were first exposed to a gray screen for 30 minutes, followed by a 30-minute presentation of mating videos (Fig. 1e). For light/dark stimuli, the screens alternated between white (RGB: 255, 255, 255) and black (0, 0, 0) at 60-minute intervals (Fig. 1j,o). During white-screen presentations, the luminance at the animal’s eye position was measured at 199.3 ± 0.7 lux, while black-screen presentations had a luminance of 3.0 ± 0.0 lux (LX-1330B lux meter, Tondaj, China).

To investigate neuronal visual responses, the visual stimuli used were full-field square wave gratings with a spatial frequency of 0.16 cycles per degree. The grating was displayed on a large screen (130° × 140°) positioned 50 cm from the viewer’s right eye using an EPSON CB-535W projector. The grating field’s horizontal and vertical meridians were rotated by 38° to match the pigeon’s natural viewing conditions ^58,59^. The luminance of black and white stripes of grating was 0.1 and 6.6 cd/m^2^, respectively. The grating moved at a speed of 8°/s for 300 ms in either the temporal-nasal (T-N) or nasal-temporal (N-T) direction ^19^. The direction of grating motion was customized for individual neuron depending on whether the motion could evoke higher neuronal visual responses. Grating motions lasting 300 ms were randomly introduced with inter-trial intervals ranging from 6 to 10 seconds. These motion stimuli were designed to probe neuronal responses while preventing the induction of saccadic or optokinetic nystagmus (OKN) eye movements (Extended Data Fig. 4).

### Pharmacological injections

To administer muscimol into the raphe complex, a Drummond™ PCR micropipette (1 to 10 µL, Thermo Fisher Scientific, USA) for microinjection using a PE-21 system (Narishige, Japan) and filled with muscimol (2%, PhytoLab, Germany). Accurate targeting of the raphe complex was achieved by initially inserting a glass-insulated tungsten microelectrode into the target area according to stereotaxic coordinates (from the interaural midpoint, P 0.25 mm, L 0.00 mm). The raphe complex was identified by the presence of well-characterized omnipause neurons, which exhibit a high tonic discharge rate that ceases during saccades in any direction ^19,23,60^. Next, the injection micropipette was connected to a gas-pressure microinjection system (Parker Picospritzer III, USA) and advanced into the raphe complex along the same trajectory as the microelectrode. Muscimol was administered in volumes of 20-50 nL. Following injections, pigeons exhibited cessation of oscillatory saccades within approximately 1 minute following the injection (Fig. 5f), with normal saccadic activity resuming after roughly 1.5 hours.

A custom-developed delivery system was for the intraocular injections. This system included a sterilized 34-gauge needle connected to a microinjection pump (Zibo Guanjie Electronic Technology, China) via a 1-mL syringe. The microinjection pump was controlled automatically using Spike2 software (version 8.24, Cambridge Electronic Design Limited, UK), enabling precise programming of injection timing and speed. The needle was carefully inserted into the eyeball through a surgically drilled window on the orbital bone, ensuring its tip was positioned in the intraocular fluid. During distinct pharmacological experiments, the following substances were injected intraocularly: D-glucose (1 mol/L, 2 μL, Sinopharm Chemical, China), L-glucose (1 mol/L, 2 μL, Sigma-Aldrich, USA), saline (2 μL, CSPC PHARMA, China), and Bay-876 (100 μmol/L, 1 μL, MedChemExpress, USA).

### Data acquisition

#### Electrooculogram recording (EOG)

An electrooculogram system was employed to monitor eye movements ^19,23,61,62^. Two EOG electrodes were carefully implanted into the anterior and posterior regions of the orbital arch, while a third electrode was positioned on the occipital bone as a reference. The eye movement signals were band-pass filtered with a range of 0.1 to 500 Hz (A-M Systems, Microelectrode AC Amplifier Model 1800, USA). Subsequently, the signals were sampled at a rate of 1000 Hz and stored for offline analysis (CED Power1401, Cambridge Electronic Design Limited, UK).

#### Assessments of glucose levels

In this study, assessments of glucose levels involve measuring the concentration of glucose in intraocular fluid and interstitial fluid of pectoral muscles, via the Continuous Glucose Monitoring (CGM) systems (Freestyle Libre 2 CGM System, Abbott, USA; GS1 CGM System, SIBIONICS, China) and the glucometer (Accu-Chek Active, Roche, Switzerland). The CGM was conducted using commercially available glucose detection probes to monitor glucose concentrations in the eye and body over hours. The probe was inserted into the eyeball through the surgical window in the orbital bone, landing in the intraocular fluid. The transmitter was secured to the skull using a UV-curing adhesive (ergo 8500 Metal, Kisling AG, Switzerland). A small amount of medical petroleum jelly (Lircon, China) was used to seal the exposed area. A second glucose monitoring probe was inserted into the pectoral muscle and fastened by adhesive tape. These probes are equipped with flexible micro-detection heads that prevent damage to animals’ tissue and have a minimal impact on their eye mobility.

We assessed the temporal sensitivity and accuracy of CGM systems to detect changes in glucose concentration. To evaluate temporal sensitivity, a higher concentration glucose solution (1ml, 15mmol/L) was introduced into the testing glucose environment (20ml, 5mmol/L) 20 seconds prior to sampling. Subsequent samples were analyzed to determine the system’s capability to detect glucose fluctuations. The response was quantified using normalized readings, calculated as CGM change divided by actual concentration change (Extended Data Fig. 2a). For accuracy evaluation, CGM measurements were validated against reference values obtained from a glumeter across glucose concentrations ranging from 2.5 mmol/L to 20 mmol/L, in increments of 2.5 mmol/L (Extended Data Fig. 2b).

#### Extracellular recordings

The single-cell recordings utilized custom-made glass-insulated tungsten microelectrodes, which had impedances ranging from 1 to 3 MΩ. The microelectrodes were laterally inserted into the left tectum to specifically target the TeO (A 5.00-6.00) and the nLM (A 5.50-6.00, L 4.00; H 5.50-6.00), and vertically introduced through the left telencephalon to target the nOPT (A 6.5-7.0, L 2.8-3.2, H 7.0). The neuronal signals underwent filtration using a band-pass filter with a range of 300 to 5,000 Hz (A-M Systems, Microelectrode AC Amplifier Model 1800, USA). These signals were then sampled at a rate of 25 kHz (CED Power1401, Cambridge Electronic Design Limited, UK). The process of spike sorting was conducted offline using the Spike2 software (Cambridge Electronic Design Limited, UK).

A photocell (6.5 cm × 6.5 cm, Shenzhen Kobry Technology, China) was attached to the screen. It has the capability to directly capture the temporal information of visual stimuli displayed on the screen at a sampling rate of 1,000 Hz by Spike2 software. The signal was simultaneously recorded with both behavioral and neuronal responses, allowing for precise data analysis.

#### Saccade video recordings

Saccadic eye movements were assessed in five avian species: pigeon, budgerigar, chukar, chicken, and duck. Birds were observed in a resting state under standard room-light conditions. A high-speed camera (120Hz, HONOR Magic6 Pro, China) equipped with an adjustable macro lens (focal length: 30 to 70 mm, MECORIGHT, China) was used to record oscillatory saccades.

#### Retinal tissue preparation and imaging

A pigeon and a chukar were randomly selected for postmortem retinal imaging. Animals were euthanized by the injection of sodium pentobarbital (200 mg/kg) into the pectoral muscles. After euthanasia, the eyes were enucleated using ophthalmic scissors. Then, a sagittal incision was made along the equator of each eyeball using a sterile scalpel blade to expose the retinal tissue. Images were acquired using the same HONOR Magic6 Pro camera under controlled illumination to minimize glare.

### Data analysis

#### Oscillatory saccade

A typical saccadic eye movement in birds consists of several oscillations occurring at a frequency of approximately 18 to 44 Hz. By employing a customized MATLAB algorithm, we successfully detected the distinctive characteristic of EOG signals. Subsequently, a manual verification process was conducted to confirm the accuracy of the results. This allowed us to precisely determine the onset and offset of each saccade, as well as the time at which the peak values occurred throughout each cycle of saccadic oscillations. The frequency of saccades and oscillations was calculated based on the number of events within the related analysis windows. Principal Component Analyses were employed to extract the primary component, which represented the saccade and oscillation frequencies.

#### Glucose levels

The CGM system continuously monitored animals’ glucose concentrations. To align with the data of saccade and oscillation, glucose levels were interpolated using a linear method between two consecutive sampled data points. Glucose levels were quantified by averaging data from three or more repetitions per animal during the gray/video and light/dark experiments. To ensure consistency, the values were normalized to the mean glucose level observed during the 30-minute baseline period preceding the video, transition of the screen or the administration of muscimol.

#### Correlation between oscillatory saccades and intraocular glucose levels

To explore the temporal relationship between eye movements and intraocular glucose, saccade frequency and oscillation frequency were computed at 1-minute time window and normalized to the gray screen baseline, matching with the intraocular glucose sampling rate. Data were averaged across trials for each animal and then pooled across individuals. During the experiment of viewing 30-minute conspecific social mating videos, scenes depicting social interactions were diverse and fragmented, accompanied by dynamically varying eye movements. A moving lag analysis was employed, applying time lags ranging from −10 to 10 minutes in 1-minute increments between saccadic eye movements and intraocular glucose levels. Pearson correlation coefficients were calculated at each lag to quantify the temporal association between oscillatory saccades and intraocular glucose concentrations (Extended Data Fig. 3). During the experiment involving viewing non-social light/dark screen switching, visual stimuli were consistent and uniform, accompanied by steady changes in eye movements. Cross-correlation analyses were performed between intraocular glucose levels and both saccade frequency and oscillation frequency, using data from −10 to 10 minutes relative to light/dark transitions to identify corresponding temporal relationships.

#### Neuronal visual responses

We successfully identified firing activity in 127 neurons from three retinorecipient structures in the visual pathways of birds. Neuronal firing rates were computed using a classic measurement approach ^63^ and averaged in 50-ms bins. In order to eliminate the effects of perisaccadic modulations from our data analyses, we initially computed the neuronal responses during saccades that were aligned with the onset and offset of saccade, respectively. The durations of saccadic inhibition and enhancement in neurons were in agreement with our prior study ^23^. Therefore, any data that fell within the time frame of perisaccadic modulations were excluded from our subsequent analysis.

In this study, a 300 ms grating motion served as a probe to assess neuronal visual responses. We calculated the average spontaneous firing rates during saccade intervals without probes, aligned to the onset of the prior saccade. For each probe, the mean spontaneous firing rate within a 300-ms window, aligned to the probe timing relative to the prior saccade, was used as the baseline. Visual responses to the grating motion were then defined by subtracting this baseline. (Fig. 2, Fig. 3, and Fig. 4). To assess the relationship between visual responses and the prior saccade, we calculated the distribution of time intervals between the two events, excluded the 5% extreme values and divided the remaining intervals equally into three stages. We then compared neuronal visual responses across these stages, as well as before and after micro-injections (Fig. 3, and Fig. 4). In the muscimol administration experiment, saccades were suppressed, and the average firing rates within a 100 ms time interval preceding the grating motion were calculated as a baseline for comparing neuronal visual responses before and after muscimol administration (Fig. 5).

#### Statistical analysis

A two-sided Wilcoxon signed-rank test was utilized to assess behavioral and neuronal visual responses for paired data, while an independent Wilcoxon rank-sum test was applied for independent observations. Visual responses across the three stages following saccades were compared using a one-way analysis of variance (ANOVA), with the Holm-Bonferroni correction applied to control for multiple comparisons. Statistical analysis was conducted with a significance level of 0.05.

## Acknowledgments

We thank Shu-Rong Wang (Institute of Biophysics), Sheng He (Institute of Biophysics), Guangwei Si (Institute of Biophysics), and members of our laboratory for their advising and commenting on the manuscript, Zhenyu Gao (Erasmus MC) for helpful discussions. Yanhui Fu (Institute of Biophysics), and Haiyan Liu (Institute of Biophysics) for their invaluable technical assistance.

## Funding

STI2030-Major Projects 2021ZD0203800

STI2030-Major Projects 2022ZD0204800

Chinese Academy of Sciences Key Program of Frontier Sciences grant QYZDB-SSW-SMC019

National Natural Science Foundation of China 31722025

National Natural Science Foundation of China 32070987

Strategic Priority Research Program of Chinese Academy of Sciences (XDB37030303)

## Author contributions

Conceptualization: YY

Methodology: YY, XX, TX, YBC

Investigation: XX, TX, YBC, LJS, CW, QW

Visualization: XX, TX, YBC, YY

Funding acquisition: YY

Project administration: YY

Supervision: YY

Writing – original draft: YY, XX

Writing – review & editing: YY, XX, TX, YBC, TZ

## Data and materials availability

Datasets supporting the findings of this study are available from the corresponding author on reasonable request.

