## supplemental fig 1-5 for "Saccades orchestrate intraocular glucose to shape visual responses in birds"

### Supplementary figures

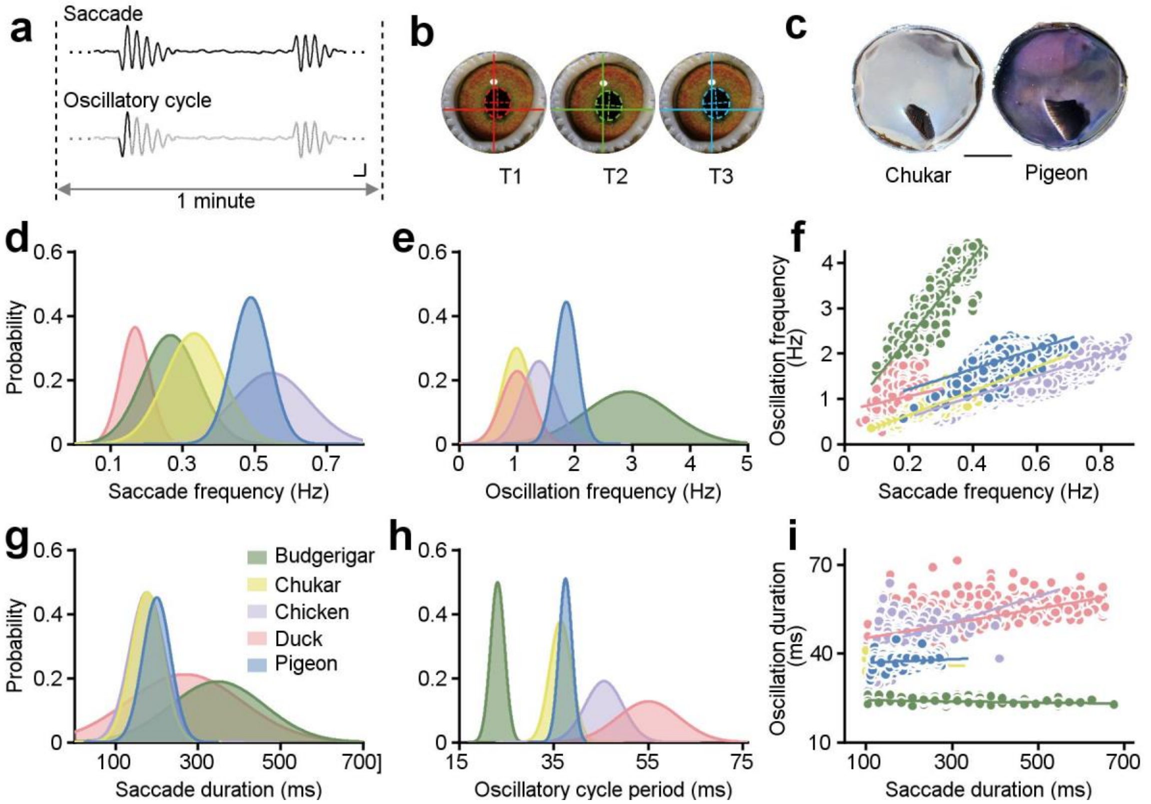


**Extended Data Fig. 1 | Comparative properties of oscillatory saccades among five avian species.**

**a,** Each saccade in birds consists of multiple oscillatory cycles. **b**, Three consecutive frames of a saccadic eye movement in a pigeon, using a high-speed camera at 120 frames per second. Solid lines represent fixed coordinate axes, while dashed lines indicate the pupil position in each frame. **c,** Images of the avian retina with the pecten oculi, illustrated using a chukar and a pigeon as examples. Scale bar: 5 mm. **d,e,** Probability density functions illustrating saccade frequency (**d**) and oscillation frequency (**e**) across five avian species during resting states under standard room-light conditions. Saccade frequency and oscillation frequency are calculated by counting the number of events occurring within each 1-minute analysis window (example shown in **a**). **f,** Relationship between saccade frequency and oscillation frequency. **g,h,** Probability density functions illustrating saccade duration (**g**), oscillation duration (**h**) across five avian species. **i,** Relationship between saccade duration and oscillation duration. The colored lines represent linear regression.

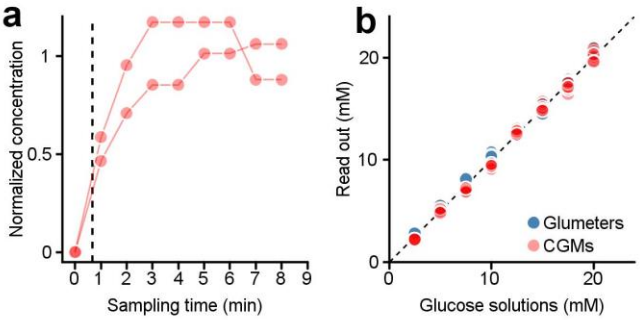


**Extended Data Fig. 2 | The temporal sensitivity and accuracy of CGM systems.**

**a,** Temporal sensitivity of CGM to glucose fluctuations assessed using two CGM sensors. A higher concentration glucose solution (15mmol/L) was introduced into the testing glucose environment (5mmol/L) 20 seconds prior to sampling (dashed line). CGM responses were quantified using normalized readings, calculated as CGM change divided by theoretical concentration change. **b,** Accuracy of CGM systems for measuring glucose concentrations. Reading out of a glucometer (blue dots) and CGM systems (red dots) were obtained using standard glucose solutions ranging from 2.5 mmol/L to 20 mmol/L, in increments of 2.5 mmol/L.

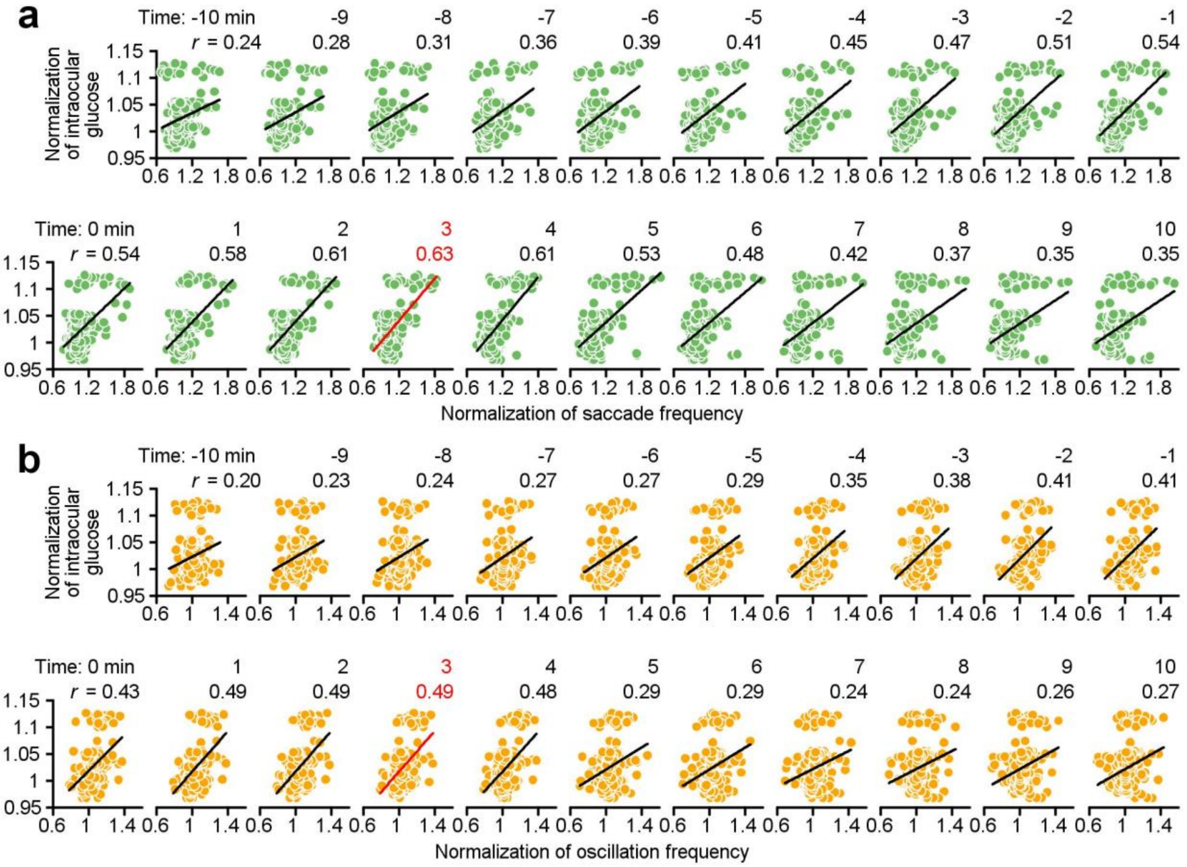


**Extended Data Fig. 3 | Correlation between oscillatory saccades and intraocular glucose** viewing 30-minute conspecific social mating videos**.**

**a,b,** Scatter plots illustrating correlations between normalized intraocular glucose levels and saccade frequency (**a**) or oscillation frequency (**b**) across time lags ranging from -10 to 10 minutes. Eye movements and intraocular glucose data were averaged across trials per animal and normalized to their gray screen baselines. Colored dots represent data from four animals. Each scatter plot corresponds to a specific time lag, with positive time lags indicating that eye movements precede intraocular glucose changes. Solid lines represent linear regression fits, with Pearson's correlation coefficient (r) shown. Red numbers highlight the time lag with highest correlation.

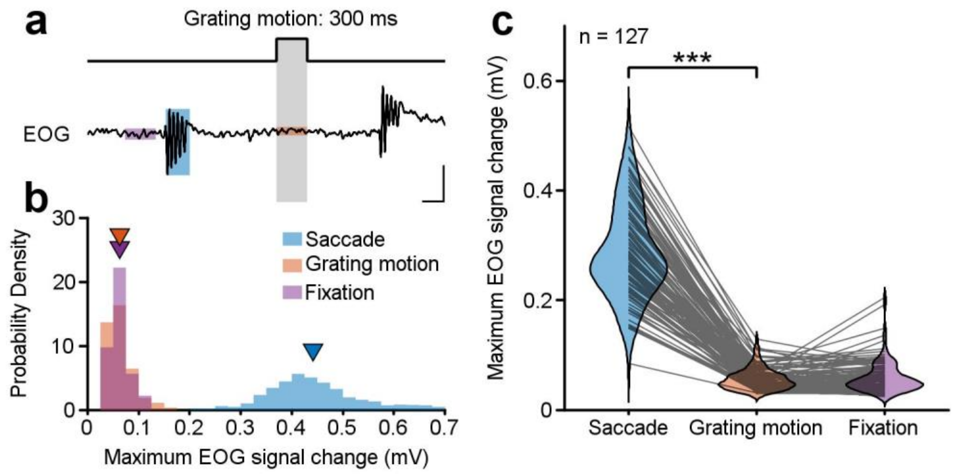


**Extended Data Fig. 4 | The 300-ms grating motion stimulus does not elicit significant eye movements in pigeons.**

**a,** Example raw EOG traces around grating motion. Three 300-ms time windows are highlighted in blue, orange, and lilac shadings, representing the saccade period, the grating motion period, and the fixation period (400 ms before the next saccade), respectively. Scale bars: 200 ms, 0.2 mV. **b,** Distribution of maximum EOG signal changes within each time window for an example neuron, with mean values marked by inverted triangles. **c,** Comparison of these EOG signals during three time windows across all 127 recorded neurons. Gray lines represent data from individual neurons. *** $P < 0.001$, Two-sided Wilcoxon signed-rank test.

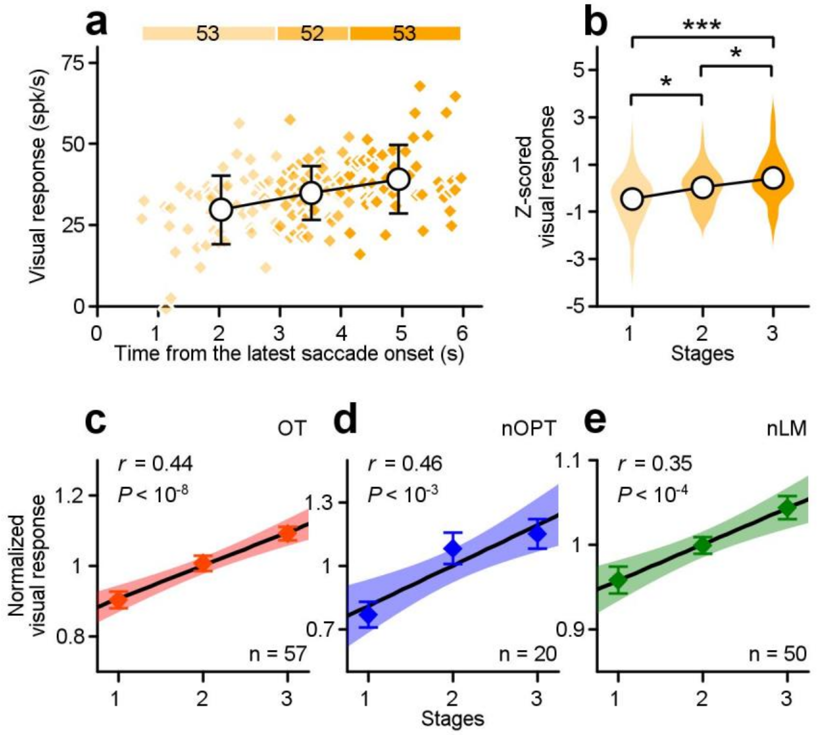


**Extended Data Fig. 5 | Consistent dynamic changes in visual responses following the latest oscillatory saccade, with stages categorized by an equal number of grating motions.**
**a,** Visual responses to each 300-ms grating motion stimulus are categorized into three stages, each containing approximately an equal number of stimuli, based on the time since the prior saccade onset. Black circles and error bars represent the mean ± SD for each stage. **b,** Comparison of the distribution of z-scored visual responses across the three stages from the example neuron (one-way ANOVA, $F_{(2,155)} = 12.08$, $P = 1.33 \times 10^{-5}$). The width of each violin reflects the density of data points, with black circles indicating the mean responses. **c-e,** Normalized visual responses across the three stages for neurons in OT (**c**), nOPT (**d**), and nLM (**e**). Data points and error bars represent mean ± SEM. Black lines represent the linear regression, while shadings are the 95% confidence interval of fitting. * $P < 0.05$, ** $P < 0.01$, *** $P < 0.001$, Two-sided Wilcoxon rank-sum test.
